# TaxonTableTools - A comprehensive, platform-independent graphical user interface software to explore and visualise DNA metabarcoding data

**DOI:** 10.1101/2020.08.24.264317

**Authors:** Till-Hendrik Macher, Arne J. Beermann, Florian Leese

## Abstract

DNA metabarcoding is increasingly used in research and application to assess biodiversity. Powerful analysis software exists to process raw data. However, when it comes to the translation of sequence read data into biological information many end users with limited bioinformatic expertise struggle with the downstream analysis and explore data only to a minor extent. Thus, there is a growing need for easy-to-use, graphical user interface (GUI) analysis software to analyse and visualise DNA metabarcoding data. We here present TaxonTableTools (TTT), a new platform independent GUI software that aims to fill this gap by providing simple and reproducible analysis and visualisation workflows. TTT uses a so-called “TaXon table” as input. This format can easily be generated within TTT from two input files: a read table and a taxonomy table that can be obtained by various published metabarcoding pipelines. TTT analysis and visualisation modules include e.g. Venn diagrams to compare taxon overlap among replicates, samples or among different analysis methods. It analyses and visualises basic statistics such as read proportion per taxon as well as more sophisticated visualisation such as interactive Krona charts for taxonomic data exploration. Various ecological analyses such as alpha or beta diversity estimates, and rarefaction analysis ordination plots can be produced directly. Data can be explored also in formats required by traditional taxonomy-based analyses of regulatory bioassessment programs. TTT comes with a manual and tutorial, is free and publicly available through GitHub (https://github.com/TillMacher/TaxonTableTools) and the Python package index (https://pypi.org/project/taxontabletools/).

## Introduction

DNA metabarcoding is increasingly used to assess biodiversity of marine (Aylagas, Borja, Muxika & Rodríguez-Ezpeleta, 2018; Zaiko, Pochon, Garcia-Vazquez, Olenin & Wood, 2018), limnic (Elbrecht, Peinert & Leese, 2017; Bush et al., 2020) and terrestrial ecosystems (Beng et al., 2016; Porter et al. 2019). It can be applied to a bulk sample containing multiple species (e.g. Elbrecht et al. 2017; Elbrecht & Steinke, 2019) or be applied to environmental DNA (eDNA; Deiner et al., 2016, Zinger et al., 2019) and allows for rapid and cost-efficient assessments of taxonomic composition. While many different DNA metabarcoding laboratory protocols have been established in recent years (see Leese et al., 2018), all DNA metabarcoding approaches are united by the amplification of a genetic marker using primers for the target group. The marker choice depends on the target taxa. Commonly used genetic markers for animals are the mitochondrial cytochrome oxidase I (COI; Leray et al., 2013, Macher, Vivancos, Piggott, Centeno, Matthaei & Leese, 2018) and different fragments of the small and large subunits of the nuclear ribosomal RNA, like the 12S (Miya et al., 2015; Hänfling et al., 2016) and 16S marker (Clarke, Soubrier, Weyrich & Cooper, 2014, Elbrecht et al., 2016). Established barcodes for plants are the matK gene (CBOL Plant Working Group et al., 2009), the large subunit of ribulose-1,5-bisphosphate carboxylase-oxygenase gene (rbcL; Hollingsworth, 2011), the internal transcribed marker (ITS; Hollingsworth, 2011, China Plant BOL Group et al., 2011), and the trnL P6 loop marker (Fahner, Shokralla, Baird & Hajibabaei, 2016), while the commonly used gene for fungi is the internal transcribed marker (ITS; Mello, Napoli, Murat, Morin, Marceddu & Bonfante, 2011; Blaalid, Kumar, Nilsson, Abarenkov, Krik & Kauserud, 2013). Following PCRs, amplicons are sequenced on high-throughput sequencer platforms and translated into digital characters. After sequencing, the obtained sequences are bioinformatically processed. For this task a number of programs such as JAMP (http://github.com/VascoElbrecht/JAMP), DADA2 (Callahan, McMurdie, Rosen, Han, Johnson & Holmes, 2016) and OBITOOLS (Boyer, Mercier, Bonin, Bras, Taberlet & Coissac, 2015) exist. While these pipelines are solely command-line based, recent efforts have been made to feature graphical user interfaces (GUIs) like for q2studio in QIIME2 (Bolyen et al., 2019) and the web-based applications SLIM (Dufresne, Lejzerowicz, Perret-Gentil, Pawlowski & Cordier, 2019) and mBRAVE (Ratnasingham, 2019). Generally, all these pipelines follow a similar workflow of processing steps, including quality control, primer trimming, quality trimming and clustering. The obtained sequences are compared to a reference database for taxonomic assignment. The largest publicly usable database for COI data is the Barcode of Life Datasystems (BOLD) database. ITS sequences are deposited in the UNITE database (Nilsson et al., 2018), while GenBank (NCBI) holds the largest repository of sequences from various markers and organisms, yet with limited curation and control. Reference sequences from online databases are often downloaded for subsequent local taxonomic assignment e.g. using the blast+ software (Camacho, Coulouris, Avagyan, Ma, Papadopoulos, Bealer & Madden, 2008). Furthermore, several tools have been published to conduct taxonomic assignment of sequences automatically against an online database, for example DADA2, JAMP, ecotag (OBITOOLS pipeline) or BOLDigger (Buchner & Leese, 2020). The resulting taxonomic assignment tables are the final outcome of all bioinformatic pipelines and as that the basis for all downstream analyses in biodiversity research or biomonitoring. For each step in the DNA metabarcoding workflow, various authors have published both laboratory protocols and bioinformatics tools. Nevertheless, until now only a few tools have been published to tackle a comprehensive downstream analysis and visualization of metabarcoding results. QIIME2 and DADA2 both include tools or instructions on how to filter tables and how to calculate and visualize e.g. diversity measurements and ordinations analyses. Another widely distributed tool is the R package ‘vegan’ (Oksanen et al., 2012), a community ecology package for ordination analyses and diversity measurements and dissimilarities. However, for the moment all these applications are mainly command line-based and require basic bioinformatics skills to be used. Methodological advancements open up the possibility of upscaling DNA metabarcoding to hundreds of samples per week and provide comprehensive investigations for large scaled biomonitoring programs. As a direct consequence, analyses of this growing amount of produced data and its translation into biological meaningful results increasingly becomes the bottleneck, which limits the uptake of the methods by non-experts and bioinformatics beginners. However, it is the biologists that need to work with the data and interpret these.

To address the clear need of analysing increased amounts of data in a user-friendly way, we developed the software *TaxonTableTools* (*TTT* in the following). TTT was developed as part of the GeDNA project, which tests the implementation of eDNA metabarcoding as part of regulatory biomonitoring. The program provides easy-to-use tools for biologists and non-bioinformaticians to analyse and visualize their metabarcoding data quickly and reproducibly via a GUI. It unites commonly used data processing steps for metabarcoding data with a set of modules used for taxonomic exploration of the results, ecological analyses as well as options to use the data as part of regulatory biomonitoring applications.

## Implementation

TaxonTableTools is written in python and available at GitHub (https://github.com/TillMacher/TaxonTableTools). Python is currently supported by all three major operating systems Windows, MacOs and Linux-based distributions (e.g. Ubuntu). Program installation only requires minimum user input. When python and pip are properly installed, the required python packages can be easily installed via pip. To improve user-friendliness, TTT comes with a mouse-driven graphical user interface (GUI), which allows the user to easily execute the various modules as well as a detailed manual and a tutorial with a test data set (figure 1).

**Figure 1:**
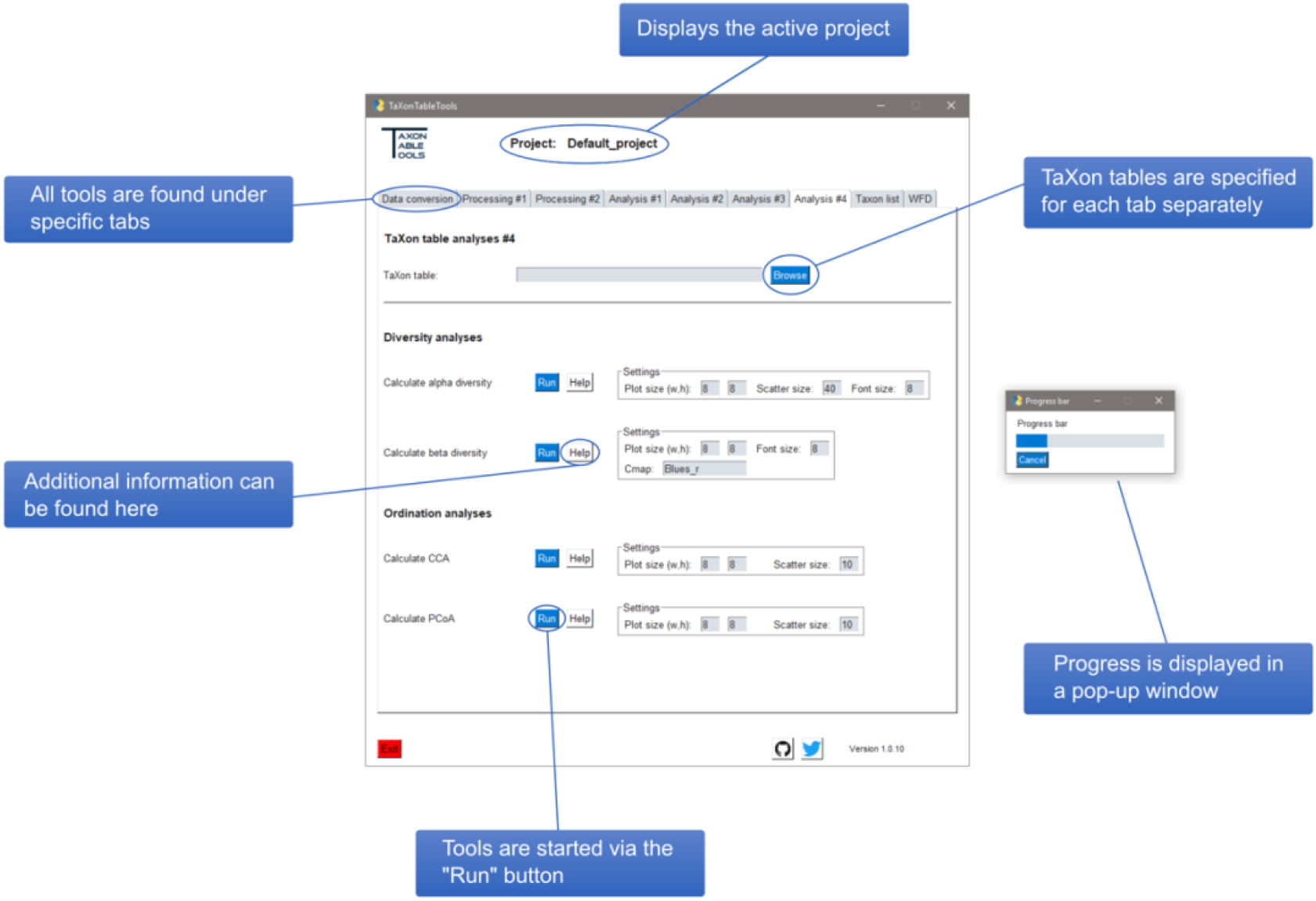
Graphical user interface of TaxonTableTools

A key advantage of TTT is a comprehensive data management structure. New projects are created within a dedicated project folder. All generated files are stored in the respective project directory, which drastically increases clarity and structure when working with different data sets or projects. When launched TTT will ask to either create a new project folder or load an already existing one. This circumvents the explicit naming of output files. Newly generated files are named according to their input file and the conducted application.

A major goal of TTT was to offer a rapid and easy tool to visualize the data for reports or publications. Thus, the standard output format for most plots is the pdf format, which retains the vectors in graphics. This allows post-processing of plots created with TTT with any vector manipulation software.

## Input formats and data conversion

### Input format requirements

TTT requires two input files, a read table and a taxonomy table. Read tables are generally referred to as a data frame, which contains the read abundances for each OTU (operational taxonomic unit) or ASV (amplicon sequence variant) per sample and its respective sequence. Read tables are generated by various published DNA metabarcoding pipelines and wrappers, e.g. JAMP, DADA2, QIIME2 or OBITOOLS. Since the output layout differs between pipelines, TTT requires a specific input layout. This can easily be created from the various other formats. Mostly only the header requires minor adjustments. Taxonomy tables are defined as a data frame that holds taxonomic information for each OTU of the read table. The layout and informational content often drastically differ, as there is no current consensus on a standard format. Taxonomy tables can for example be created using QIIME2, SLIM, blast+, BOLDigger or even be compiled manually. As the standard input format TTT uses the output format from the BOLDigger tool. As a requirement, the same OTUs have to be present in both the taxonomy table from BOLDigger and the respective read table. Information on phylum, class, order, family, genus and species level are required and intermediate classifications are not allowed. Only one hit per OTU is accepted, which requires a prior filtering step. This can be performed with BOLDigger. The range of accepted input formats regarding the read and taxonomy tables will be subject of change in future versions of TTT to reduce user-based reformatting prior to the import in TTT.

### Getting started: Merging of read and taxonomy tables

Initially TTT requires two tables: a) a read table and b) a taxonomy table. Both tables are merged into one table that contains information of both individual formats. Thus, one main table is created, which is referred to as “TaXon table” and is the standard input for all downstream applications in TTT. TaXon tables can also be manually created. The format can automatically be checked to prevent downstream errors. The main advantage of the TaXon table format is the combination of all relevant data for DNA metabarcoding derived taxonomic tables into one file. Thus, for the sake of simplicity, the redundancy of two separate tables is solved, and subsequently data can be more rapidly accessed. For example, read distributions per OTU and sample can be directly investigated, while immediately checking the OTU’s taxonomy and if necessary, also its respective sequence. This is advantageous for both manually investigations and bioinformatic processing.

### TaXon table processing

#### Replicate processing

Most data sets require some processing steps prior to analyses. Here, TTT offers various tools to merge, filter and convert TaXon tables. All processing tools are optional and can be used as separate modules. The processing workflow generally starts with the processing of sample replicates. Replicates are generally recommended in DNA metabarcoding analyses to tackle PCR and sequencing stochasticity (e.g. Weigand & Macher, 2018; Mata, Rebelo, Amori, McCracken, Jarman & Beja, 2018). No specific replicate design is required. Replicates are recognized via the sample names, which have to be marked with a trailing underscore and a user-defined symbol at the end (e.g. commonly used “_rep1”, “_rep2” or “_a”, “_b”). Here, the first module allows to filter OTUs by keeping only OTUs that are present in all replicates of one sample. In detail, this will exclude OTUs that are not present in all replicates of one sample, by setting the OTU read counts to zero. When the research interest is focused on low abundant or rare OTUs, this module is not recommended, since it might lead to exclude real, but rare OTUs. Afterwards replicates can be merged into one representative sample, by calculating the sum of reads for each OTU of the replicates. This will drastically reduce redundancy in the TaXon table and is often useful for downstream analyses that do not require separate replicates.

#### Adding metadata

Metadata are generally used as criterion to infer differences between samples by assigning specific values according to location, environmental factors (e.g. ph or salinity) or classifications (e.g. European Water Framework Directive ecological status). Sample-specific metadata can then be used for downstream analyses. For example, differences in taxa distributions can be investigated (site occupancy) or statistical analyses can be performed using the metadata to explore the sample set with the principle coordinate analysis (PCoA) or canonical-correlation analysis (CCA). TTT allows the user to create a separate metadata table that is automatically linked with the respective TaXon table. Metadata categories can be manually assigned as columns, where the rows stand for the respective value of the sample. Currently only the inclusion of metadata for samples is implemented. However, the TTT roadmap includes the analyses of metadata for OTUs in a future version.

#### Conversion to incidence data

The use of read abundances as a proxy for specimen counts or biomass estimates has been subject of discussion with the development of DNA metabarcoding. Due to PCR stochasticity, varying primer binding efficiency and sequencing bias, there is often only a weak correlation between read abundances and specimen counts or biomass (Elbrecht & Leese, 2015; Elbrecht et al., 2017, Bista et al., 2018), although for several cases with improved primer settings exceptions exist (e.g. Schenk, Geisen, Kleinboelting & Traunspurger, 2019). Thus, often it is recommended to convert the read abundance data to incidence data for biodiversity analyses. However, this conversion comes with a downside, since incidence data limits the pool of appropriate diversity estimate analyses.

### TaXon table analysis

#### Getting first insights

To get a first overview of the data set, it is helpful to visualize the number of reads, number of OTUs and OTUs assigned to species level in a plot (figure 2a). This plot allows investigations of the overall quality of the data set. Generally, negative controls should represent only a fraction of the overall reads. Furthermore, samples that have drastically lower read counts and thus often also less OTUs and OTUs on species level, should be considered to be removed from the data set. They are often prone to create outliers in statistical analyses or alter the perspective of between locations or categories comparisons.

**Figure 2:**
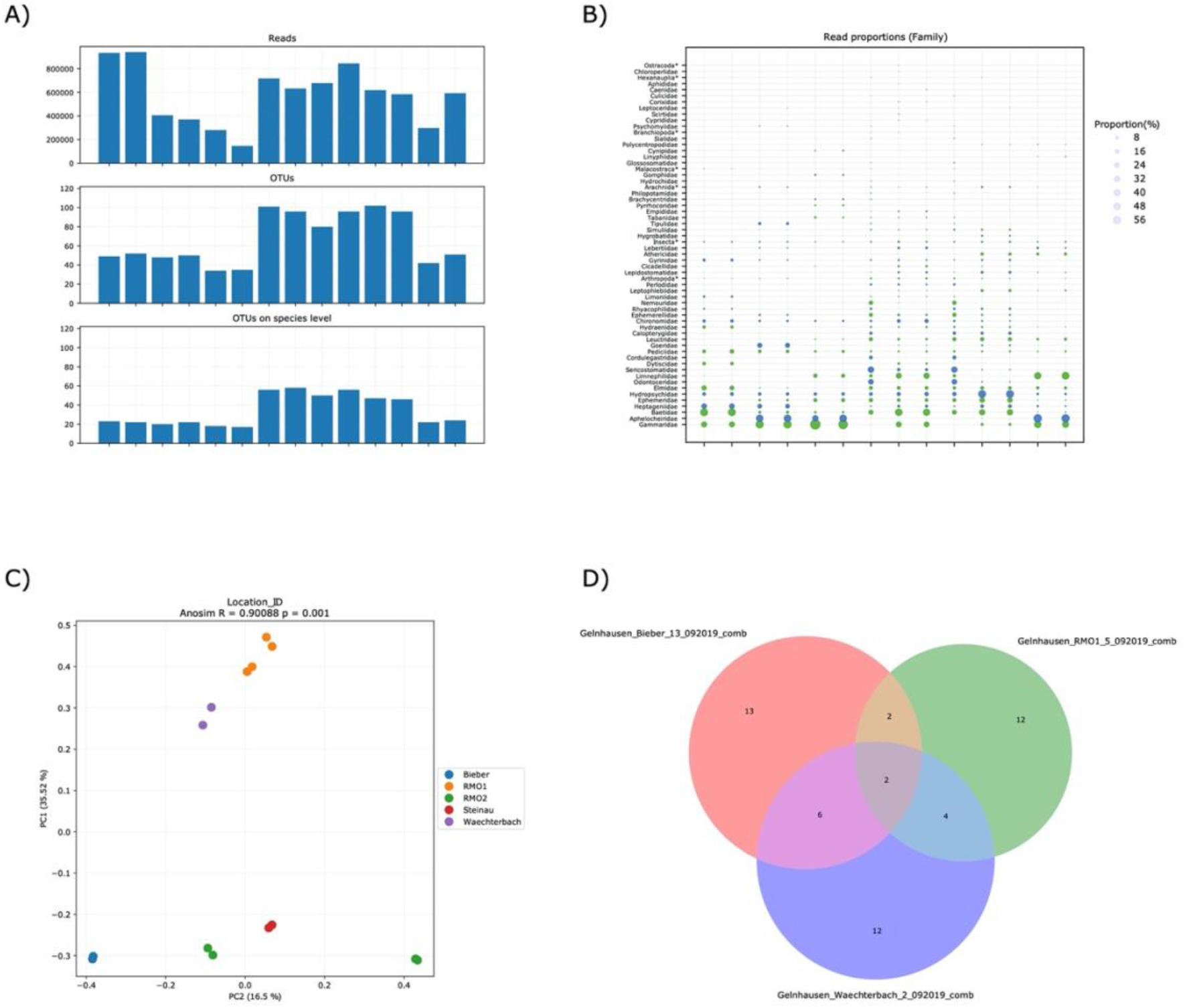
Exemplary graphical output produced with TaxonTableTools: Basis statistics give a first overview of the data set by plotting the number of reads, number of OTUs and number of OTUs on species level for all samples (A). Read proportions can be plotted in a scatter plot, where the circles size represents the proportion of the respective taxon in the sample (B). Sample names on the x-axis have been removed for sub-plots A and B. Correlations between samples can be investigated by performing a principle coordination analysis (PCoA), which is based on Jaccard distances (C). An analysis of similarities (ANOSIM) and p-test is performed automatically. Taxa overlaps of up to three samples can be visualized with Venn diagrams (D).

#### Read proportions

Read proportions can be illustrated in a scatter plot. Contrary to the commonly applied illustration in a bar plot, a scatter plot remains readable even for larger data sets (figure 2b). Each taxon is represented by its own entity so that the dependency on colors schemes is no longer required.

#### Taxonomic richness and resolution

Measuring the taxonomic richness of a sample assemblage is an essential objective in every biodiversity analysis. In classical ecology terms, the species richness is defined as the number of species in an ecological community, landscape or region. The most straightforward computation of species richness is to count the number of OTUs or species in the data set. The species richness can either be calculated for the whole data set or for each sample itself.

However, identification to species level is often not possible. Many species remain undescribed and there is a lack of reference sequences for a vast number of species. Still, higher taxonomic levels (i.e. genus or family) can hold information for assessing biodiversity. The overall taxonomic resolution of a data set can be visualized in a bar chart, which plots the number of OTUs assigned to the respective level as lowermost rank. The taxonomic resolution can be used as an indicator for several potential sources of bias, like varying primer binding efficiency or bioinformatics process bias (e.g. remaining primer sequences). These can act as sources for a reduced taxonomic resolution, as OTUs are often not assigned to species level in consequence.

#### Rarefaction

Sample-based rarefaction is a commonly applied method to infer if the number of samples taken was sufficient to capture the complete species richness. This method computes the number of species when samples are drawn at random without replacement from a set of samples. Replicating the drawing results (e.g. 1000 times) substantially enhances the robustness of rarefaction and allows the calculation of standard deviations. Sample-based rarefaction curves can be calculated using the Rarefaction curve tool. Nevertheless, when interpreting rarefaction curves, the methodical limitations of DNA metabarcoding must be considered.

#### OTU abundance pie charts and Krona charts

OTU proportions per taxon can be investigated via pie charts. Here, individual pie charts are created for every taxonomic level in the TaXon table. This method is independent from read abundances. It can be used to display the OTU distributions on higher taxonomic levels. Furthermore, potential clustering bias can be inferred by investigating the OTU proportions on lower taxonomic levels, when e.g. looking for cryptic species. On the other hand, Krona tools allows the hierarchical illustration of read proportions (Beermann, Werner, Elbrecht, Zizka & Leese 2020, Ondo, Bergman & Phillippy, 2011). OTUs that have been assigned to the same taxon (e.g. one species) are merged. The data can be explored by zooming through a multi-layered, interactive pie-chart that can be viewed with any modern web browser.

#### Comparing samples via Venn diagrams

The taxon composition between up to three TaXon tables can be displayed via venn diagrams (figure 2d). The comparison is performed on each taxonomic level in the TaXon table. To allow for more detailed investigations, the groupings are written to a separate excel file, while only the number of taxa in each group of the venn diagram is displayed. It is not recommended to draw venn diagrams of more than three data sets, as the plot quickly becomes confusing.

#### Diversity analyses and ordination methods

DNA metabarcoding data is often used for diversity analyses. TTT offers calculation of alpha and beta diversity and ordination analyses. The implemented tools are mostly dependent on the python package scikit-bio (http://scikit-bio.org/). All diversity analyses require an incidence data TaXon table. The alpha diversity calculation is based on the number of OTUs per sample, which are displayed as a scatter plot. Beta diversity is calculated as Jaccard-distances, which are illustrated in a distance matrix. Furthermore, a Jaccard-distance based principle coordinate analysis (PCoA) can be performed (figure 2c). A canonical-correlation analysis (CCA) tool is also implemented. For both ordination analyses, it is possible to choose two axes from all available axes for plotting.

#### Report and taxa list

A taxa list can be created from the TaXon table. This table includes all found taxa from the input TaXon table and reduces redundant hits. Optionally, for each hit that was identified on species level, a link to the Global Biodiversity Information Facility (GBIF) website is created. The GBIF database (https://www.gbif.org/) is accessed via the application programming interface (API). These links allow for quick investigations of the taxon list, particularly for checking unfamiliar taxa. Furthermore, statistics can be calculated for each taxon. These include the absolute number of reads per taxon and the relative proportion within the data set, the occupancy across all samples, the number of OTUs identified as the respective taxon and the intraspecific distances for taxa with multiple OTUs. In addition to the taxon list, a report file is created. This report file includes many relevant information on how the data was processed from the wet lab to bioinformatics processing. This information can be filled out in the GUI and enhances the data documentation in hindsight.

#### Conversion to regulatory assessment programs

One additional aim of TTT is to convert metabarcoding data sets into formats required by tools used for regulatory frameworks such as the European Water Framework Directive (WFD, Directive 2000/60/EC). Monitoring activities for the WFD, and for counterparts in other areas and for other ecosystems, aim to provide standardized assessments of the ecological quality of waterbodies derived from biota. In the initial version, we provide the opportunity to convert metabarcoding lists into a format that can be used as input to the German Water Framework Directive analysis tool. This online tool (www.gewaesser-bewertung-berechnung.de) is designed to allow the upload of taxa lists from monitoring activities, from which the ecological quality according to the German river assessment scheme is calculated. In addition, many supporting metrics (such as feeding types or habitat preferences of macroinvertebrates) are calculated. Upload requires a species-station table in an Excel or ASCII format with species in rows and stations in columns, giving the abundance (or alternatively the presence / absence) of the recorded taxa. Each taxon is accompanied by an ID that allows for linking the taxon to its specific autecological characteristics. For comparability reasons, the system standardizes the taxonomy to an operational taxa list, which defines for each taxon the taxonomic level achievable by identification in routine water management. As an alternative to the direct upload, the system offers a batch mode, allowing large data sets to be automatically read from databases of water authorities and the assessment results to be returned. TTT provides species station tables in the format required by this system, including the taxon ID, that can be directly uploaded and used for river assessment.

## Verification data set

For verification of TaxonTableTools a test data set was chosen, which was generated during the advanced master module “Molecular Ecology” of the University of Duisburg-Essen. Samples were taken in September 2019 in the Rhein-Main-Observatorium (RMO). Kick-net samples of macrozoobenthic invertebrates were taken at five different sites in the RMO. Samples were sorted in the field, collecting up to 300 specimens per site and stored in ethanol. Samples were then laid out over night for drying in petri dishes. Afterwards, samples were transferred into Turrax tubes and pulverized with two IKA Ultra-Turrax devices. DNA was extracted using salt precipitation as described in Weiss and Leese (2016). The amplification was performed using a two-step PCR with the BF2/BR2 primer set (see Elbrecht & Leese, 2017 for more details), pooled and sequenced on a MiSeq v2 2×250 bp. The retrieved sequences were demultiplexed and processed using the JAMP pipeline v.067. The taxonomic assignment was performed with BOLDigger (pre-release version).

## Acknowledgments

We thank the leeselab members, especially Dominik Buchner for comments and feedback on the program. We thank Daniel Hering (University of Duisburg-Essen, Aquatic Ecology) for input and helpful contributions to the MS. We thank Jan Koschorreck and Jens Arle (German Federal Environment Agency) for their feedback on the manuscript. The software was developed as part of the GeDNA project funded by the German Federal Environmental Agency (FKZ 3719 24 2040).

## Data Accessibility

Project name: “TaxonTableTools: A versatile, platform independent tool for reproducible analyses and visualization of DNA metabarcoding data”; Project home page: https://github.com/TillMacher/TaxonTableTools; Operating system(s): Platform independent; Programming language: Python 3; Other requirements: Krona tools (https://github.com/marbl/Krona); License: MIT licence; Any restrictions to use by nonacademics: No restrictions.

## Author Contributions

T.M. conceived and designed the study and wrote the Python package. A.B. and F.L. provided input to the package and supervised the project. T.M., F.L. and A.B. wrote the paper. All authors read and approved the final manuscript.

## References

Aylagas, E., Borja, Á., Muxika, I., & Rodríguez-Ezpeleta, N. (2018). Adapting metabarcoding-based benthic biomonitoring into routine marine ecological status assessment networks. Ecological Indicators, 95, 194–202. doi: 10.1016/j.ecolind.2018.07.044

Beermann, A. J., Werner, M. T., Elbrecht, V., Zizka, V. M. A., Leese, F. (2020). DNA metabarcoding improves the detection of multiple stressor responses of stream invertebrates to increased salinity, fine sediment deposition and reduced flow velocity. Science of the Total Environment (accepted)

Beng, K. C., Tomlinson, K. W., Shen, X. H., Surget-Groba, Y., Hughes, A. C., Corlett, R. T., & Slik, J. W. F. (2016). The utility of DNA metabarcoding for studying the response of arthropod diversity and composition to land-use change in the tropics. Scientific Reports, 6(1), 24965. doi: 10.1038/srep24965

Bista, I., Carvalho, G. R., Tang, M., Walsh, K., Zhou, X., Hajibabaei, M., … Creer, S. (2018). Performance of amplicon and shotgun sequencing for accurate biomass estimation in invertebrate community samples. Molecular Ecology Resources, 18(5), 1020–1034. doi: 10.1111/1755-0998.12888

Blaalid, R., Kumar, S., Nilsson, R. H., Abarenkov, K., Kirk, P. M., & Kauserud, H. (2013). ITS1 versus ITS2 as DNA metabarcodes for fungi. Molecular Ecology Resources, 13(2), 218–224. doi: 10.1111/1755-0998.12065

Bolyen, E., Rideout, J. R., Dillon, M. R., Bokulich, N. A., Abnet, C. C., Al-Ghalith, G. A., … Caporaso, J. G. (2019). Reproducible, interactive, scalable and extensible microbiome data science using QIIME 2. Nature Biotechnology, 37(8), 852–857. doi: 10.1038/s41587-019-0209-9

Boyer, F., Mercier, C., Bonin, A., Bras, Y. L., Taberlet, P., & Coissac, E. (2016). obitools: a unix-inspired software package for DNA metabarcoding. Molecular Ecology Resources, 16(1), 176–182. doi: 10.1111/1755-0998.12428

Buchner, D., & Leese, F. (2020). BOLDigger – a Python package to identify and organise sequences with the Barcode of Life Data systems. Metabarcoding and Metagenomics, 4, e53535. doi: 10.3897/mbmg.4.53535

Bush, A., Monk, W. A., Compson, Z. G., Peters, D. L., Porter, T. M., Shokralla, S., … Baird, D. J. (2020). DNA metabarcoding reveals metacommunity dynamics in a threatened boreal wetland wilderness. Proceedings of the National Academy of Sciences, 117(15), 8539–8545. doi: 10.1073/pnas.1918741117

Callahan, B. J., McMurdie, P. J., Rosen, M. J., Han, A. W., Johnson, A. J. A., & Holmes, S. P. (2016). DADA2: High-resolution sample inference from Illumina amplicon data. Nature Methods, 13(7), 581–583. doi: 10.1038/nmeth.3869

Camacho, C., Coulouris, G., Avagyan, V., Ma, N., Papadopoulos, J., Bealer, K., & Madden, T. L. (2009). BLAST+: architecture and applications. BMC Bioinformatics, 10, 421. doi: 10.1186/1471-2105-10-421

CBOL Plant Working Group, Hollingsworth, P. M., Forrest, L. L., Spouge, J. L., Hajibabaei, M., Ratnasingham, S., … Little, D. P. (2009). A DNA barcode for land plants. Proceedings of the National Academy of Sciences, 106(31), 12794–12797. doi: 10.1073/pnas.0905845106

China Plant BOL Group, Li, D.-Z., Gao, L.-M., Li, H.-T., Wang, H., Ge, X.-J., … Duan, G.-W. (2011). Comparative analysis of a large dataset indicates that internal transcribed spacer (ITS) should be incorporated into the core barcode for seed plants. Proceedings of the National Academy of Sciences, 108(49), 19641–19646. doi: 10.1073/pnas.1104551108

Clarke, L. J., Soubrier, J., Weyrich, L. S., & Cooper, A. (2014). Environmental metabarcodes for insects: in silico PCR reveals potential for taxonomic bias. Molecular Ecology Resources, 14(6), 1160–1170. doi: 10.1111/1755-0998.12265

Deiner, K., Bik, H. M., Mächler, E., Seymour, M., Lacoursière-Roussel, A., Altermatt, F., … Bernatchez, L. (2017). Environmental DNA metabarcoding: Transforming how we survey animal and plant communities. Molecular Ecology, 26(21), 5872–5895. doi: 10.1111/mec.14350

Dufresne, Y., Lejzerowicz, F., Perret-Gentil, L. A., Pawlowski, J., & Cordier, T. (2019). SLIM: a flexible web application for the reproducible processing of environmental DNA metabarcoding data. BMC Bioinformatics, 20(1), 88. doi: 10.1186/s12859-019-2663-2

Elbrecht, V., & Leese, F. (2015). Can DNA-Based Ecosystem Assessments Quantify Species Abundance? Testing Primer Bias and Biomass—Sequence Relationships with an Innovative Metabarcoding Protocol. PLOS ONE, 10(7), e0130324. doi: 10.1371/journal.pone.0130324

Elbrecht, V., & Leese, F. (2017). Validation and Development of COI Metabarcoding Primers for Freshwater Macroinvertebrate Bioassessment. Frontiers in Environmental Science, 5. doi: 10.3389/fenvs.2017.00011

Elbrecht, V., Peinert, B., & Leese, F. (2017). Sorting things out: Assessing effects of unequal specimen biomass on DNA metabarcoding. Ecology and Evolution, 7(17), 6918–6926. doi: 10.1002/ece3.3192

Elbrecht, V., & Steinke, D. (2019). Scaling up DNA metabarcoding for freshwater macrozoobenthos monitoring. Freshwater Biology, 64(2), 380–387. doi: 10.1111/fwb.13220

Elbrecht, V., Taberlet, P., Dejean, T., Valentini, A., Usseglio-Polatera, P., Beisel, J.-N., … Leese, F. (2016). Testing the potential of a ribosomal 16S marker for DNA metabarcoding of insects. PeerJ, 4, e1966. doi: 10.7717/peerj.1966

Fahner, N. A., Shokralla, S., Baird, D. J., & Hajibabaei, M. (2016). Large-Scale Monitoring of Plants through Environmental DNA Metabarcoding of Soil: Recovery, Resolution, and Annotation of Four DNA Markers. PLOS ONE, 11(6), e0157505. doi: 10.1371/journal.pone.0157505

Hänfling, B., Handley, L. L., Read, D. S., Hahn, C., Li, J., Nichols, P., … Winfield, I. J. (2016). Environmental DNA metabarcoding of lake fish communities reflects long-term data from established survey methods. Molecular Ecology, 25(13), 3101–3119. doi: 10.1111/mec.13660

Hollingsworth, P. M. (2011). Refining the DNA barcode for land plants. Proceedings of the National Academy of Sciences, 108(49), 19451–19452. doi: 10.1073/pnas.1116812108

Jari Oksanen and F. Guillaume Blanchet and Michael Friendly and Roeland Kindt and Pierre Legendre and Dan McGlinn and Peter R. Minchin and R. B. O’Hara and Gavin L. Simpson and Peter Solymos and M. Henry H. Stevens and Eduard Szoecs and Helene Wagner. (2019). vegan: Community Ecology Package.

Leese, F., Bouchez, A., Abarenkov, K., Altermatt, F., Borja, Á., Bruce, K., … Weigand, A. M. (2018). Chapter Two - Why We Need Sustainable Networks Bridging Countries, Disciplines, Cultures and Generations for Aquatic Biomonitoring 2.0: A Perspective Derived From the DNAqua-Net COST Action. In D. A. Bohan, A. J. Dumbrell, G. Woodward, & M. Jackson (Eds.), Advances in Ecological Research (pp. 63–99). doi: 10.1016/bs.aecr.2018.01.001

Leray, M., Yang, J. Y., Meyer, C. P., Mills, S. C., Agudelo, N., Ranwez, V., … Machida, R. J. (2013). A new versatile primer set targeting a short fragment of the mitochondrial COI region for metabarcoding metazoan diversity: application for characterizing coral reef fish gut contents. Frontiers in Zoology, 10(1), 34. doi: 10.1186/1742-9994-10-34

Macher, J.-N., Vivancos, A., Piggott, J. J., Centeno, F. C., Matthaei, C. D., & Leese, F. (2018). Comparison of environmental DNA and bulk-sample metabarcoding using highly degenerate cytochrome c oxidase I primers. Molecular Ecology Resources, 18(6), 1456–1468. doi: 10.1111/1755-0998.12940

Mata, V. A., Rebelo, H., Amorim, F., McCracken, G. F., Jarman, S., & Beja, P. (2019). How much is enough? Effects of technical and biological replication on metabarcoding dietary analysis. Molecular Ecology, 28(2), 165–175. doi: 10.1111/mec.14779

Mello, A., Napoli, C., Murat, C., Morin, E., Marceddu, G., & Bonfante, P. (2011). ITS-1 versus ITS-2 pyrosequencing: a comparison of fungal populations in truffle grounds. Mycologia, 103(6), 1184–1193. doi: 10.3852/11-027

Miya, M., Sato, Y., Fukunaga, T., Sado, T., Poulsen, J. Y., Sato, K., … Iwasaki, W. (2015). MiFish, a set of universal PCR primers for metabarcoding environmental DNA from fishes: detection of more than 230 subtropical marine species. Royal Society Open Science, 2(7), 150088. doi: 10.1098/rsos.150088

Nilsson, R. H., Larsson, K.-H., Taylor, A. F. S., Bengtsson-Palme, J., Jeppesen, T. S., Schigel, D., … Abarenkov, K. (2019). The UNITE database for molecular identification of fungi: handling dark taxa and parallel taxonomic classifications. Nucleic Acids Research, 47(D1), D259–D264. doi: 10.1093/nar/gky1022

Oksanen, J., Guillaume Blanchet, F., Friendly, M., Kindt, R., Legendre, P., McGlinn, D., … Wagner, H. (2019). vegan: Community Ecology Package.

Ondov, B. D., Bergman, N. H., & Phillippy, A. M. (2011). Interactive metagenomic visualization in a Web browser. BMC Bioinformatics, 12(1), 385. doi: 10.1186/1471-2105-12-385

Porter, T. M., Morris, D. M., Basiliko, N., Hajibabaei, M., Doucet, D., Bowman, S., … Venier, L. (2019). Variations in terrestrial arthropod DNA metabarcoding methods recovers robust beta diversity but variable richness and site indicators. Scientific Reports, 9(1), 18218. doi: 10.1038/s41598-019-54532-0

Ratnasingham, S. (2019). mBRAVE: The Multiplex Barcode Research And Visualization Environment. Biodiversity Information Science and Standards, 3, e37986. doi: 10.3897/biss.3.37986

Schenk, J., Geisen, S., Kleinboelting, N., & Traunspurger, W. (2019). Metabarcoding data allow for reliable biomass estimates in the most abundant animals on earth. Metabarcoding and Metagenomics, 3, e46704. doi: 10.3897/mbmg.3.46704

Weigand, A. M., & Macher, J.-N. (2018). A DNA metabarcoding protocol for hyporheic freshwater meiofauna: Evaluating highly degenerate COI primers and replication strategy. Metabarcoding and Metagenomics, 2, e26869. doi: 10.3897/mbmg.2.26869

Weiss, M., & Leese, F. (2016). Widely distributed and regionally isolated! Drivers of genetic structure in Gammarus fossarum in a human-impacted landscape. BMC Evolutionary Biology, 16(1), 153. doi: 10.1186/s12862-016-0723-z

Zaiko, A., Pochon, X., Garcia-Vazquez, E., Olenin, S., & Wood, S. A. (2018). Advantages and Limitations of Environmental DNA/RNA Tools for Marine Biosecurity: Management and Surveillance of Non-indigenous Species. Frontiers in Marine Science, 5. doi: 10.3389/fmars.2018.00322

Zinger, L., Bonin, A., Alsos, I. G., Bálint, M., Bik, H., Boyer, F., … Taberlet, P. (2019). DNA metabarcoding—Need for robust experimental designs to draw sound ecological conclusions. Molecular Ecology, 28(8), 1857–1862. doi: 10.1111/mec.15060

